# Parallel Characterization of *cis*-Regulatory Elements for Multiple Genes Using CRISPRpath

**DOI:** 10.1101/2021.02.19.431931

**Authors:** Xingjie Ren, Mengchi Wang, Bingkun Li, Kirsty Jamieson, Lina Zheng, Ian R. Jones, Bin Li, Maya Asami Takagi, Jerry Lee, Lenka Maliskova, Tsz Wai Tam, Miao Yu, Rong Hu, Lindsay Lee, Armen Abnousi, Gang Li, Yun Li, Ming Hu, Bing Ren, Wei Wang, Yin Shen

## Abstract

Current pooled CRISPR screens for *cis*-regulatory elements (CREs) can only accommodate one gene based on its expression level. Here, we describe CRISPRpath, a scalable screening strategy for parallelly characterizing CREs of genes linked to the same biological pathway and converging phenotypes. We demonstrate the ability of CRISPRpath for simultaneously identifying functional enhancers of six genes in the 6-thioguanine-induced DNA mismatch repair pathway using both CRISPR interference (CRISPRi) and CRISPR nuclease (CRISPRn) approaches. 60% of the identified enhancers are known promoters with distinct epigenomic features compared to other active promoters, including increased chromatin accessibility and interactivity. Furthermore, by imposing different levels of selection pressure, CRISPRpath can distinguish enhancers exerting strong impact on gene expression from those exerting weak impact. Our results offer a nuanced view of *cis*-regulation and demonstrate that CRISPRpath can be leveraged for understanding the complex gene regulatory program beyond transcriptional output at scale.

## Main

*Cis*-regulatory elements (CREs) are key regulators for spatial-temporal control of gene expression. Mutations in CREs can contribute to complex diseases by modulating gene expression over long genomic distances^1-3^. Thus, functionally characterizing CREs can provide important insight into gene regulation mechanisms and enable us to better interpret non-coding genetic variants associated with diseases. Despite the fact that tremendous numbers of candidate CREs have been mapped by biochemical signature^4^, our knowledge of whether, how, and how much these putative CREs are functional on gene expression remain scarce in the human genome. Pooled CRISPR screens have been developed for testing CREs in their native chromatin context by monitoring the transcriptional levels for the gene of interest^5-11^. Although results from these studies have made significant contributions to the annotation of functional DNA elements, challenges remain in pooled CRISPR screens of CREs. First, CRISPR screens for enhancers based on gene expression levels largely depend on generating reporter knock-in cell lines^7^ or using FlowFISH signals^8^. These procedures, involving generation of reporter lines and selection of cells with positive hits by flow cytometry, are time-consuming and difficult to scale up to multiple genes in the same experiment. Second, the approaches of using gene expression as the screening phenotype^9, 10^ fail to connect the functions of DNA elements from transcriptional regulation at the molecular level to interpretable cellular and physiological functions. Third, in cases of CRE screens using phenotypes such as cell proliferation and survival^11, 12^, they fail to quantify the effect sizes of enhancers on transcriptional output.

To address these limitations, we developed CRISPRpath, a pooled CRISPR screening approach to simultaneously characterizing CREs for multiple target genes involved in the same biological pathway. CRISPRpath allows us to screen functional DNA elements based on phenotypes associated with well-defined biological pathways. We demonstrate the capacity of CRISPRpath by performing CRISPR interference (CRISPRi) and nuclease (CRISPRn) screens for six genes in human induced pluripotent stem cells (iPSCs), and reveal different strengths of enhancer functions by imposing varying levels of selection pressure on the cells.

## Results

### Leveraging CRISPRpath for parallel characterization of CREs for multiple genes in iPSCs

To characterize candidate *cis*-regulatory elements (CREs) for multiple genes within the same pooled CRISPR screening, we designed and applied CRISPRpath to six genomic loci containing six genes (*HPRT1, MSH2, MSH6, MLH1, PMS2, PCNA*) involved in the 6-thioguanine (6TG)-induced mismatch repair (MMR) (**Fig. 1a**). The MMR pathway is highly conserved and essential for the maintenance of genome stability^13^. The MMR pathway recognizes DNA mismatches caused by 6TG treatment and induces cell apoptosis^14, 15^. On the other hand, cells with a malfunctioning MMR pathway, due to aberrant expression levels of 6TG metabolism genes or MMR genes, may survive during 6TG treatment. Employing the properties of the MMR pathway, we used cell survival for selecting cells with the reduced expression of MMR genes due to defects in enhancer activities (**Fig. 1b**). To design the screening library, we first identified open chromatin regions by performing Assay for Transposase Accessible Chromatin using sequencing (ATAC-Seq) in WTC11 iPSCs. We included all open chromatin regions defined by ATAC-seq peaks located 1Mb upstream and 1Mb downstream of each of the six genes (spanning a total of 10.6 Mb genomic regions) as candidate CREs for functional characterization (**Supplementary Fig. 1a, b, Supplementary Table 1**). We then designed a sgRNA library with 32,383 distal sgRNAs targeting 294 distal ATAC-seq peaks, 2,755 proximal sgRNAs targeting 81 ATAC-seq peaks overlapped with transcription start site (TSS) and coding regions of the six genes, and 625 non-targeting sgRNAs with genomic sequences in the same genomic loci but are not followed by PAM sequences (**Supplementary Fig. 1c, Supplementary Table 2**). In total, we included 35,763 sgRNAs in the library with an average of 110 sgRNA per ATAC-seq peak (**Supplementary Fig. 1d, Supplementary Fig. 2a, b**). We generated a lentiviral library expressing these sgRNAs and transduced this library into two engineered WTC11 iPSC lines, one expressing doxycycline-inducible dCas9-KRAB (CRISPRi) and the other doxycycline-inducible Cas9 (CRISPRn)^16^, both at a multiplicity of infection (MOI) of 0.5 (**Fig. 1b**).

**Figure 1.**
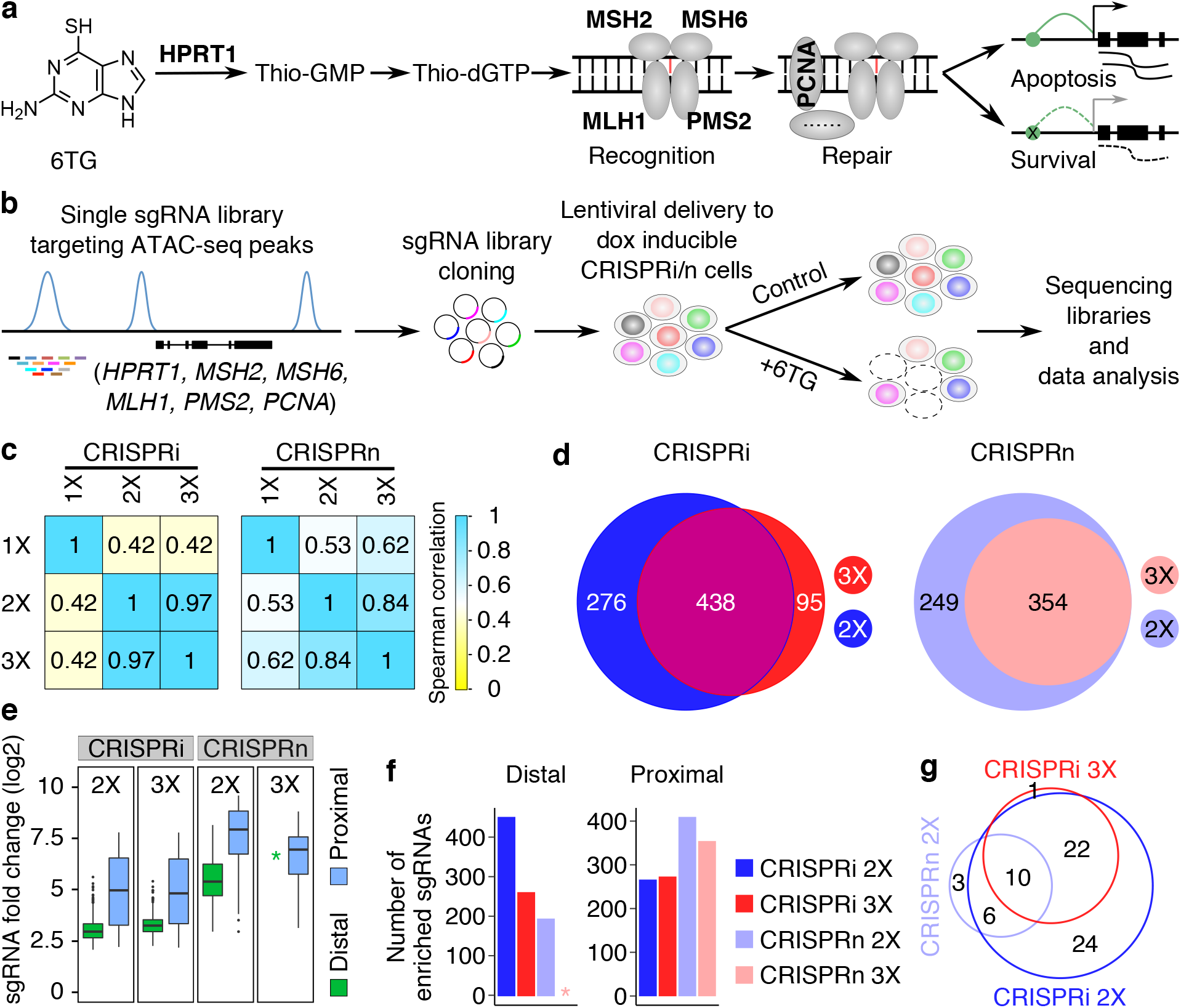
CRISPRpath for identifying enhancers of multiple genes. (**a**) Six genes (*HPRT1, MSH2, MSH6, MLH1, PMS2* and *PCNA*) in the 6TG-induced mismatch repair process were used for CRISPRpath screen in this study. (**b**) Schematic of the CRISPRpath screening strategy with 6TG treatment in iPSCs. Cell survival was used as readout for the screen. (**c**) Spearman correlation analysis of sgRNA ranking based on fold change for CRISPRpath screens with different 6TG concentrations (1X, 2X and 3X). (**d**) Venn diagram shows the overlapping enriched sgRNAs identified from the screens with 2X and 3X 6TG treatments. (**e**) Box plots show the fold change of the enriched distal and proximal sgRNAs from 2X, 3X CRISPRi and CRISPRn screens. Star indicates no enriched distal sgRNA was identified from 3X CRISPRn screen. Boxplots indicate the median, interquartile range (IQR), Q1 − 1.5 × IQR and Q3 + 1.5 × IQR. (**f**) Bar plot shows the number of enriched distal and proximal sgRNAs from 2X, 3X CRISPRi and CRISPRn screens. Star indicates no enriched distal sgRNA was identified from 3X CRISPRn screen. (**g**) Venn diagram shows the identified enhancers from each CRISPRpath screen.

To carry out the screening, we pre-determined the minimal lethal concentration of 6TG at 80 ng/mL for CRISPRi and CRISPRn iPSC lines (see **Methods** for more details), and applied three different 6TG concentrations (1X: 80 ng/mL, 2X: 160 ng/mL, 3X: 240 ng/mL) in both CRISPRi and CRISPRn screens. We extracted and sequenced DNA samples from the survival cells seven days after 6TG treatment to determine enriched sgRNAs by comparing the results to that of the control cells taken after sgRNA library infection before the 6TG treatment (**Fig. 1b**). To avoid confounding signals generated by off-target effects of low-quality sgRNAs^17^, we only used sgRNAs with high specificity (defined as specificity score > 0.2^18^, and without any off-target sites with sequence similarity of ≤2 mismatches) for data analysis. This led to the use of a total of 12,702 high-quality sgRNAs with an average of 38 sgRNAs per ATAC-seq peak for analysis (**Supplementary Fig. 1e, f, Supplementary Fig. 2c, d**). We performed each screen in two biological replicates with each pair of replicates exhibiting high reproducibility (**Supplementary Fig. 2e**). We compared the abundance of each sgRNA between the 6TG treated population and the control population using a negative binomial model, and computed the fold change and *P* value to quantify the effect size and the significance of enrichement of each sgRNA. We used the 5% percentile of the *P* values from non-targeting control sgRNAs as the empirical significance threshold to achieve a false discovery rate of 5%. sgRNAs with *P* value less than the empirical significance threshold and with fold change > 2 were defined as enriched. (**Supplementary Fig. 3)**. As expected, sgRNA targeting TSS and coding region were identified as positive hits from both CRISPRi and CRISPRn screens exhibiting greater fold change in CRISPRn screens compared to the CRISPRi screens (**Fig. 1e, Supplementary Fig. 4a**). We also observed enrichment of sgRNA bias towards coding regions over TSS regions for CRISPRn screen (**Supplementary Fig. 4b**). These results are consistent with CRISPRi functioning best near TSS by inhibiting transcription, and CRISPRn can disrupt gene function by generating indels downstream of TSS^19, 20^.

Further sgRNA fold-change ranking analysis revealed strong positive correlation between the screens with 2X and 3X 6TG treatment for both CRISPRi and CRISPRn screen (Spearman correlation, CRISPRi = 0.97, CRISPRn = 0.84) (**Fig. 1c**) with the correlations for proximal sgRNAs being higher than for distal sgRNAs (**Supplementary Fig. 4c**). On the contrary, results from the 1X screen correlated poorly with either 2X or 3X screens (**Fig. 1c**), suggesting more substantial selection pressure (2X and 3X) can reduce background noise in CRISPRpath screens. Thus, we used sgRNAs enriched from 2X and 3X screens data for identifying active enhancers in the following section (**Fig. 1d**).

### CRISPRi is more efficient than CRISPRn in pooled CRISPR screens of CREs

Performing CRISPRpath with CRISPRi and CRISPRn in the same genetic background with an identical sgRNA library offers a unique opportunity for comparing the efficacies of CRISPRi and CRISPRn in pooled CRISPR screens of CREs. We noticed that CRISPRn screens recovered fewer enriched distal sgRNAs than CRISPRi screens (**Fig. 1f**). This is possibly due to the fact that CRISPRi-mediated heterochromatin formation can more effectively perturb CREs compared to CRISPRn-mediated genetics perturbations. We then called an candidate element as an enhancer if there are at least 3 enriched sgRNAs in that CRE. Based on this criterion, we identified 62 and 33 enhancers from the 2X and 3X CRISPRi screen, respectively, and 19 enhancers from the 2X CRIPSRn screen. (**Fig. 1g, Supplementary Table 3**). However, no enhancer was identified from the 3X CRISPRn screen, indicating either the CRISPRn induced mutations did not lead to any strong effect on gene expression to make the cells survive the 3X 6TG treatment or there are insufficent numbers of sgRNAs exhibiting deleterious effects on the tested DNA elements to satisfy our criterion of calling functional enhancers. In total, 66 unique enhancers were identified for the six target genes with CRISPRpath under different 6TG treatments (**Fig. 1g**). Together, we demonstrate CRISPRpath can simultaneously identify enhancers for multiple target genes with CRISPRi outperforming CRISPRn. For the following analysis, we focused on the 63 enhancers identified from the 2X and 3X CRISPRi screens (**Fig. 1g**).

### Genomic feature of CRISPRpath identified enhancers

To determine the genomic feature of the enhancers, we plotted all the tested elements by their genomic locations and enrichment scores (average of log_2_(fold change) of enriched sgRNAs of each element) (**Fig. 2a**). Not surprisingly, our data suggest that each gene can be regulated by multiple enhancers with the identified functional enhancers having no position bias relative to the TSS. The average distance between an enhancer and its paired TSS is about 530 Kb (**Fig. 2b**) with an average of 10 interval genes between an identified enhancer and its target gene pairs (**Fig. 2c**). Interestingly, we observed a weak negative correlation between the enhancer enrichment score and the distance between an enhancer and its paired TSS (**Fig. 2d**, Pearson correlation, ρ = -0.36, *P* = 0.01), suggesting enhancers near to TSS tend to have higher regulatory activity compared to enhancers further away from their target genes. It is worth noting, the relative positions for the enriched sgRNAs exhibited no preference relative to ATAC-seq peaks (**Fig. 2e, Supplementary Fig. 5a**) and no preference for the strand on which the sgRNAs were designed (**Supplementary Fig. 5b**), consistent with our knowledge that CRISPRi mediated heterochromatin spreads over hundreds of base pairs in distance^21^.

**Figure 2.**
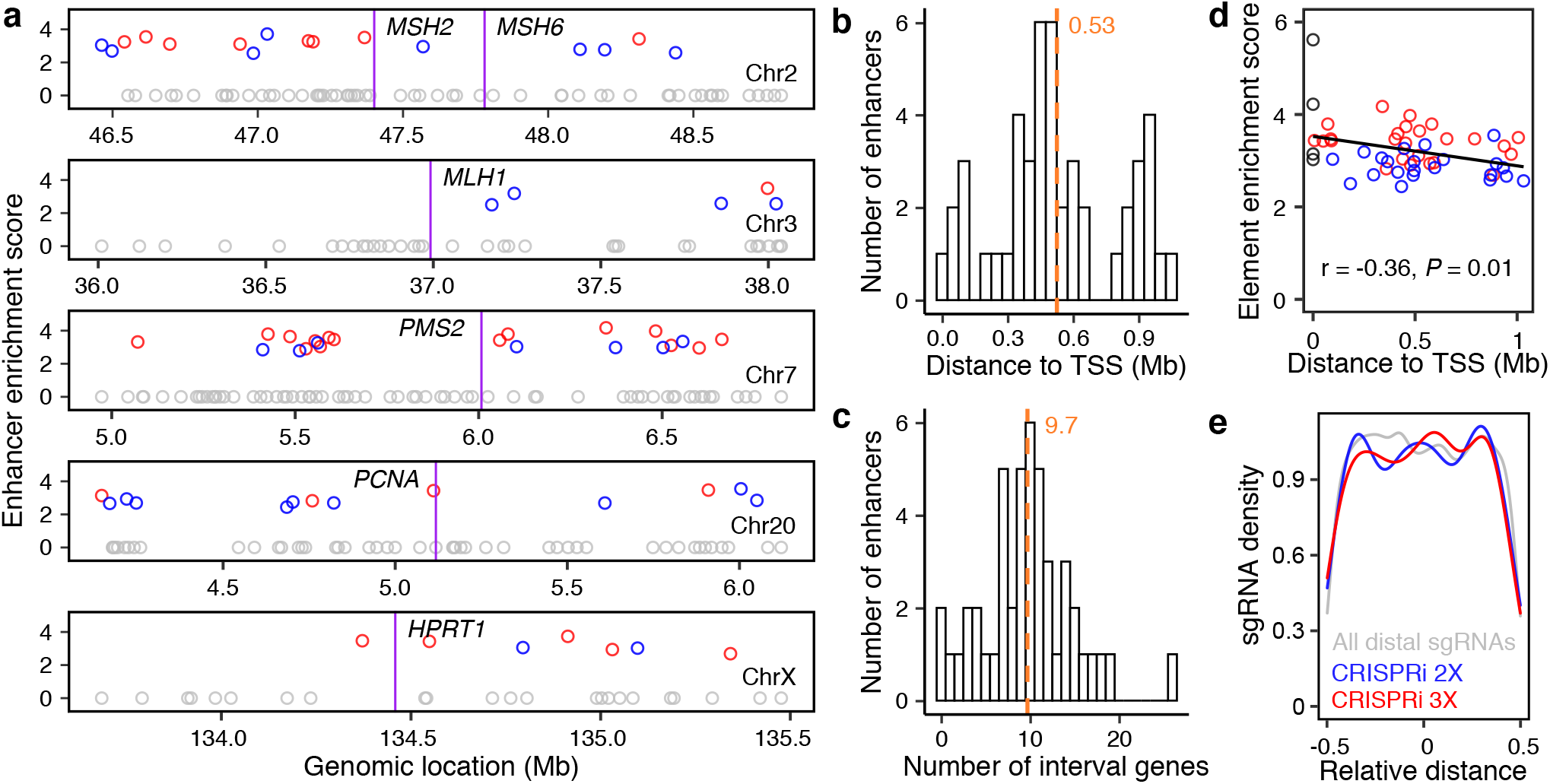
Genomic features of identified enhancers from CRISPRpath using CRISPRi. (**a**) Genomic locations of identified enhancers relative to TSS. Circles indicate enhancers identified from the CRISPRi 3X screen (red), enhancers uniquely identified from the CRISPRi 2X screen (blue), tested CREs that are not identified as enhancers (grey). Purple lines label the location of each target gene. (**b**) Histogram shows the distance distribution between identified enhancers and their paired TSS. (**c**) Histogram shows the number of interval genes between enhancers and their target gene TSS. Mean is indicated with an orange dashed line and only the enhancers for *MLH1, PMS2, PCNA, HPRT1* are included in **b** and **c**. (**d**) A weak negative correlation is observed between enrichment score and genomic distance between enhancers and their target genes (Pearson correlation, r = -0.36, *P* = 0.01). Black circles indicate promoters. The red and blue circles are enhancers showing in **a**. (**e**) Density plot shows no significant difference (two-tailed two-sample Kolmogorov–Smirnov test) for the distribution of all distal sgRNAs (gray), enriched distal sgRNAs from 2X (blue) and 3X screen (red) CRISPRi screens.

Previous studies have revealed that promoters can function as enhancers^7, 22^. Indeed, 60% (38 out of 63) of the functional enhancers identified in CRISPRi screens overlapped with annotated promoters, providing an excellent opportunity to further explore the genomic features of these enhancer-like promoters. To validate whether these promoters function as bona fide enhancers, we targeted three enhancer-like promoters with CRISPRi. We confirmed significant downregulation of their target genes including *MSH6, MSH2*, and *PCNA* (**Fig. 3a-c**). In contrast, shRNAs against the transcripts from these promoters (*SOC5, FOXN2*, and *TMEM230*) only led to a significant downregulation of its own transcripts and did not affect their target gene expression (**Fig. 3a-c**). These results confirm that these promoter sequences identified by CRISPRpath can function as enhancers. Although it has been shown that enhancer-like protomors are enriched with active chromatin marks and physically close to target genes^7^, it is not clear whether enhancer-like promoters have unqiue genome features that can differentiate them from other regular active promoters. To this end, we compared chromatin accessibility, occupancy of histone 3 lysine 4 trimethylation (H3K4me3), histone 3 lysine 27 acetylation (H3K27ac) and CTCF, transcription, and chromatin interactivity levels between enhancer-like promoters and all other active promoters that did not show enhancer activity in our CRISPRi screens. We show that enhancer-like promoters exhibit higher chromatin accessibility, higher level of transcription, stronger H3K4me3 and H3K27ac signals than those at other active promoters (**Fig. 3d**). On the other hand, we did not observe a significant difference for CTCF binding signals between enhancer-like promoters and control promoters (**Fig. 3d**). Furthermore, by evaluating chromatin interaction data using H3K4me3 Proximity Ligation-Assisted ChIP-seq (PLAC-seq), we show enhancer-like promoters have significantly more and stronger interactions compared to control promoters (**Fig. 3e**).

**Figure 3.**
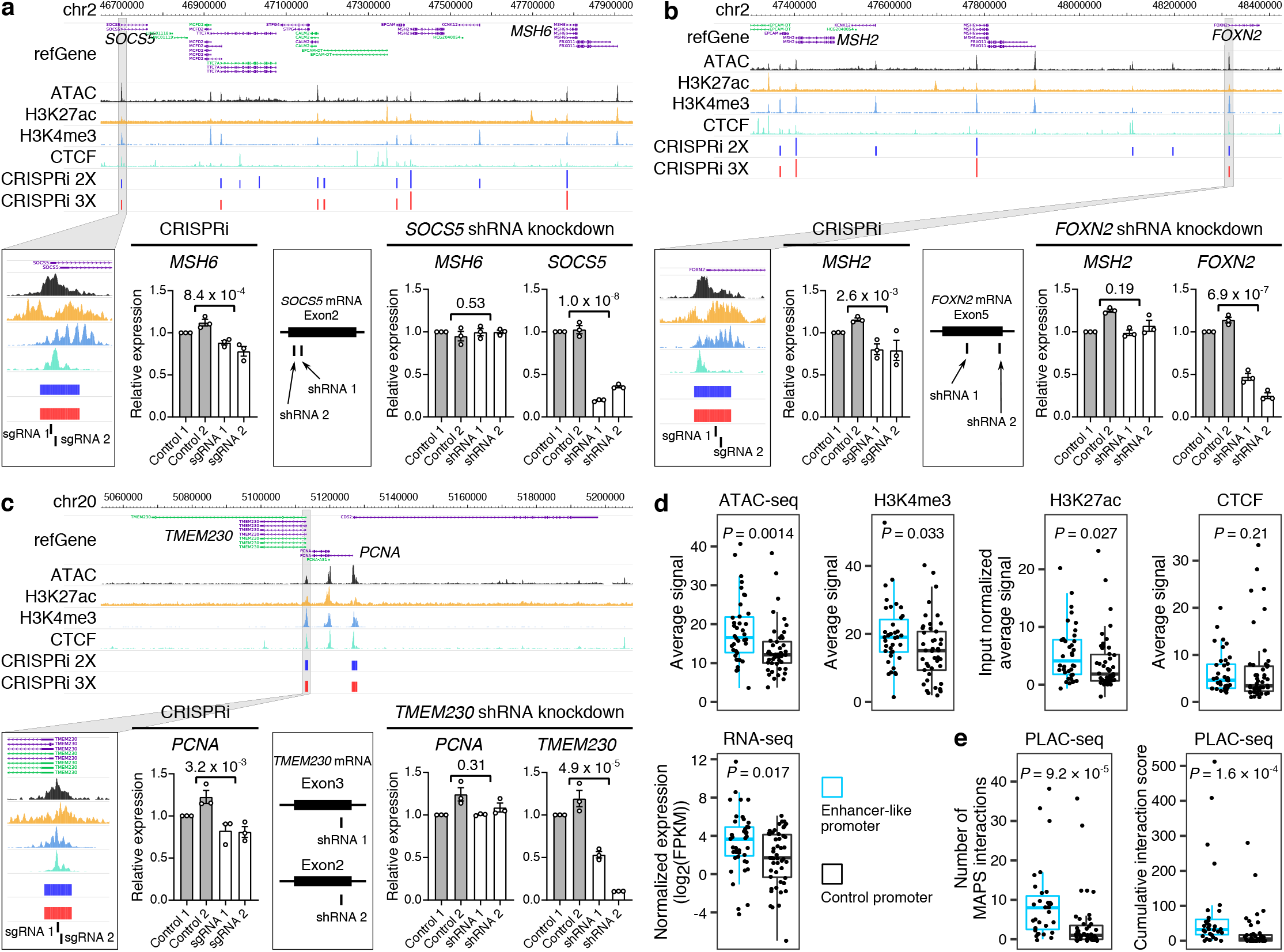
Enhancer-like promoters act as functional enhancers. (**a, b, c**) Three examples of promoters function as enhancers. CRISPRi silencing of the promoter region of *SOCS5, FOXN2* and *TMEM230* results in significant downregulation of *MSH6, MSH2* and *PCNA*, respectively. shRNA knockdown of *SOCS5, FOXN2* and *TMEM230* can only downregulate *SOCS5, FOXN2* and *TMEM230* expression. Three independent replicates per condition and two independent sgRNAs or shRNAs per replicate were used for each experiment. *P* values are from two-tailed two-sample t-test. (**d**) Average signal enrichment of ATAC-seq, gene transcription, H3K4me3, H3K27ac and CTCF binding for enhancer-like promoters (n = 38) and control promoters (n = 47). *P* values are from Wilcoxon test. Boxplots indicate the median, IQR, Q1 − 1.5 × IQR and Q3 + 1.5 × IQR. (**e**) Number of H3K4me3 mediated chromatin interactions and cumulative interaction score for enhancer-like promoters (n = 31) and control promoters (n = 43). Boxplots indicate median, IQR, Q1 − 1.5 × IQR and Q3 + 1.5 × IQR. *P* values are calculated from Wilcoxon test.

### CRISPRpath is capable of distinguishing enhancers with distinct effect sizes

Gene expression often is a result of combinatorial regulatory effects from multiple *cis*-regulatory elements^11, 23^. Understanding how individual enhancers contribute to gene expression in a quantitative manner is an important first step in dissecting how enhancers orchestrate precise transcriptional control. We seek a new strategy to differentiate enhancers based on their effect sizes on gene expression using CRISPRpath. We hypothesized that cells with drastic down-regulation of MMR genes have a fitness advantage under higher 6TG concentration than cells with modest down-regulation of MMR genes. Consistent with this hypothesis, proximal sgRNAs exhibit larger fold changes than distal sgRNAs (**Fig. 1e**) because perturbing proximal regions has more profound effects on gene down-regulation than perturbing distal regulatory regions. Based on these observations, we hypothesize that enhancers identified under different selection pressure represent distinct regulatory strengths on transcriptional activation. We noticed that enhancers identified under strong selection pressure (3X) have higher enrichment scores compared to enhancers uniquely identified under weak selection pressure (2X), with the TSS regions manifesting the highest enrichment scores (**Fig. 4a**). Thus, enhancers identified in the 3X screen are strong enhancers (n=33), while enhancers uniquely identified in the 2X screen are weak enhancers (n=30) (**Fig. 1g**). To confirm the quantitative effect of enhancers on target gene expression, we tested 11 strong and 10 weak enhancers using CRISPRi followed by RT-qPCR measurement of the corresponding target gene expression (**Fig. 4b, Supplementary Fig. 6a**). We show perturbations of strong enhancers led to significantly more down-regulation of target gene expression (mean down-regulation of target gene by 21%) than perturbations of weak enhancers (mean down-regulation of target gene by 6%), with the perturbations of TSS regions achieving the strongest down-regulation of target genes, by an average of 68% reduction in gene expression (**Fig. 4b, Supplementary Fig. 6a, b**). These quantitative effects on target gene expression are consistent with the enrichment scores from our CRISPRpath screens (**Supplementary Fig. 6c**) and demonstrate the capacity of distinguishing enhancers with different effect sizes by imposing different levels of selection pressures.

**Figure 4.**
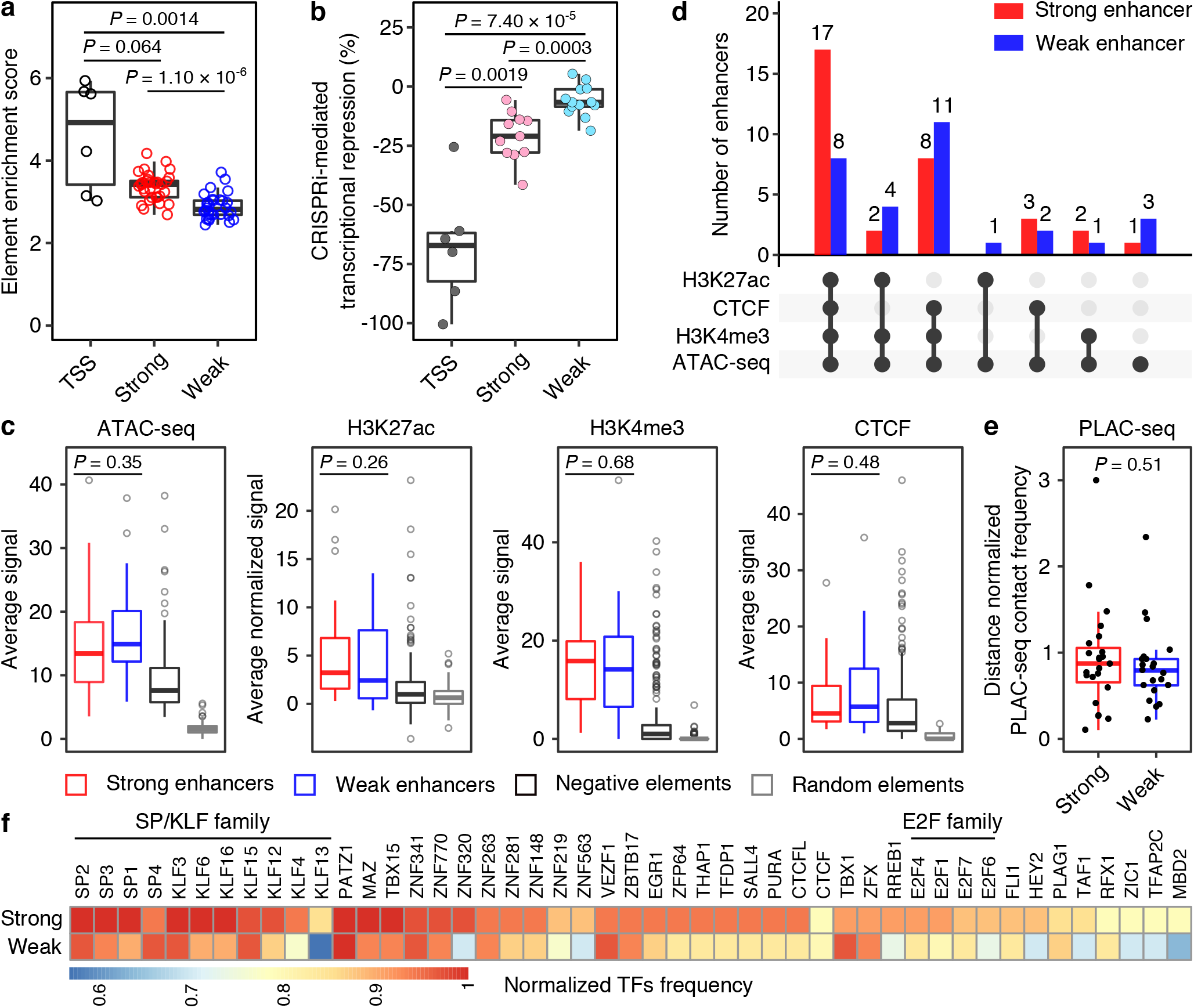
CRISPRpath can distinguish weak and strong enhancers by imposing different selection pressures. (**a**) Box plots show the enrichment score of the tested elements. TSS regions (black circles) show highest enrichment scores. Enhancers uniquely identified from the lower selection pressure (CRISPRi 2X, blue circles) exhibit lower enrichment scores compared to the enhancers identified from the higher selection pressure (CRISPRi 3X, red circles). *P* values are from Wilcoxon test. (**b**) Box plots show the CRISPRi perturbation at enhancers induced various degrees of transcriptional repression of target genes measured with RT-qPCR. Each dot represents the average value from three biological replicates. CRISPRi targeting TSS regions (dark gray) achieved the highest transcriptional repression. CRISPRi targeting strong enhancers (pink) leads a more substantial transcription silencing on target gene compared to CRISPRi targeting weak enhancers (cyan). *P* values are from Wilcoxon test. (**c**) Enrichment analysis of ATAC-seq, H3K27ac, H3K4me3, CTCF binding signals for strong (n = 33) and weak (n = 30) enhancers. Boxplots indicate the median, IQR, Q1 − 1.5 × IQR and Q3 + 1.5 × IQR. *P* values for the difference between strong and weak enhancers are from Wilcoxon test; see Supplementary Table 7 for *P* values of all pairwise comparisons. (**d**) Intersection of genomic features for weak enhancers (blue bar) and strong enhancers (red bar). (**e**) Distance normalized H3K4me3 PLAC-seq contact frequency for strong (n = 23) and weak (n = 21) enhancers. Only the enhancers for *MLH1, PMS2, PCNA, HPRT1* are included (see Methods). Boxplots indicate the median, IQR, Q1 − 1.5 × IQR and Q3 + 1.5 × IQR. *P* value is from Wilcoxon test. (**f**) Heatmap shows the frequency of transcription factor motifs found in strong and weak enhancers.

We further explored chromatin features of strong and weak enhancers by analyzing chromatin accessibility, H3K4me3, H3K27ac, and CTCF binding signals in these regions. At individual chromatin mark level, while CRISPRpath-identified enhancers were more accessible, and enriched with active chromatin marks, such as H3K4me3 and H3K27ac, and CTCF binding compared to negative elements or random elements (**Fig. 4c**), we did not observe significant differences between strong and weak enhancers in the chromatin features we individually examined. However, strong enhancers tend to have more active chromatin signatures than weak enhancers (**Fig. 4d**), suggesting combined signatures of active chromatin can be a better indicator of enhancer strength. Strong enhancers tend to have higher distance normalized PLAC-seq contact frequencies with their target promoters than weak enhancers, though not statistically significant, possibly due to the small sample size in this study (**Fig. 4e**). We obtained similar results by expanding this analysis for characterized enhancers in K562 cells and mouse embryonic stem cells^6, 8^ (**Supplementary Fig. 7**), which reinforces the idea that enhancers with larger effects on gene expression tend to have higher chromatin interactions with their cognate promoters. To explore the possible mechanisms that drive enhancer activities in a quantitative manner, we evaluated potential transcription factors (TFs) binding motifs in strong and weak enhancer sequences. Both strong and weak enhancers are enriched with CTCF binding motif (**Fig. 4f**). Indeed, most of strong and weak enhancers are bound by CTCF (**Fig. 4d**), consistent with the notion that CTCF-mediated chromatin loops are essential for gene activation^24^. Furthermore, strong enhancers and weak enhancers have differential enrichment with TFs binding motifs. For example, the binding motifs for SP/KLF family^25^ and E2F family^26, 27^ appear more frequently in strong enhancers compared to weak enhancers, suggesting these strong enhancers could be major docking sites for master regulators in iPSCs (**Fig. 4f**).

## Discussion

CRISPR-mediated high throughput screening using bulk cells allows for the functional characterization of regulatory elements in their native genomic context. However, current approaches are limited to validating a small number of regulatory elements for a single gene^5, 7, 9, 12, 28, 29^. To overcome this bottleneck, we developed CRISPRpath, a strategy for functional characterization of enhancers for multiple genes simultaneously by leveraging the genes involved in the same biological pathway so that the effects can be measured via a define phenotype. For example, alpha-toxin resistance phenotype can be used to identify CREs for 17 genes in glycosylphosphatidylinositol (GPI)-anchor synthesis pathway^30^. CRISPRpath can also be leveraged to identify CREs for protein folding regulators that contribute to the endoplasmic reticulum stress-response pathway^31^ using UPRE reporter in mammalian cells. Since CRISPR screen technology is widely used, the CRISPRpath strategy is readily applicable to simultaneously identifying enhancers for genes converging in the defined biological processes and pathways across different cell types. Compared to the existing pooled CRISPR screens of CREs^5, 7, 8, 10-12, 28, 29, 32^, CRISPRpath is scalable with additional benefits of connecting DNA elements to cellular function, beyond the most standard molecular phenotype of gene expression.

Promoters can function as enhancers more widespread than expected, with more than half of the enhancers identified for MMR genes in our study being previously annotated promoters. This is consistent with previous reports that enhancer-like promoters are more prevalent for ubiquitously expressed genes^33^. Enhancer-like promoters are more accessible compared to other promoters, possibly because these regions are required to be more open to accommodate additional transcriptional machinery such as TF for activating target gene expression besides their own transcription^34^. Enhancer-like promoters also exhibit significantly higher levels of chromatin interactions with distal regions compared to other active promoters. This observation can be explained by the fact that enhancer-like promoters will not only form chromatin loops with their distal target genes, but also with CREs for controlling the expression of their own genes.

Genomic studies of chromatin marks have revealed hundreds of thousands candidate CREs in the human genome but with very little quantitative information regarding how CREs contribute to gene regulation^35, 36^. Using CRISPRpath, we can systematically classify enhancers based on their effect sizes on transcription. Identifying and charaterizing the effect size for each individual enhancer is the critical first step to future studies of their combinatory effects on target gene expression. Interestingly, strong and weak enhancers can not be distinguished by individual epigenetic marks we examined. One possible explanation for this observation is that chromatin features only mark enhancer’s identity but do not quantify enhancer activity. On the other hand, the strong and weak enhancers we identified may regulate other genes differently from regulating the MMR gene. Interestingly, strong enhancers tend to harbor more than one active chromatin signature, which indicates that enhancer activities are regulated by multiple epigenetic factors, for example, TF mediated transcriptional regulation. Differential TFs binding motifs observed within strong and weak enhancers suggest that enhancer strength is modulated by TF binding. Future studies that further integrating TF binding datasets with functional data of enhancers will shed light on the molecular mechanisms that drives enhancers’ effect sizes on gene regulation.

## Methods

### Cell culture

Doxycycline inducible CRISPRi and CRISPRn WTC11 iPSC lines were purchased from Gladstone Stem Cell Core. Both CRISPRi and CRISPRn WTC11 iPS cells were cultured on Matrigel-coated (Corning, 354277) plates with Essential 8™ Medium (Life Technologies, A1517001). iPSCs were passaged using Accutase (STEMCELL Technologies, 07922) and 10 µM ROCK inhibitor Y-27632 (STEMCELL Technologies, 72302). HEK293T cells were cultured in Dulbecco’s modified Eagle’s medium (Gibco, 11995065) with 10% fetal bovine serum (CPSSerum, FBS-500). HEK293T cells were passaged with Trypsin-EDTA (Gibco, 25200072). All the cells were grown with 5% CO_2_ at 37°C and verified mycoplasma free using the MycoAlert Mycoplasma Detection Kit (Lonza, LT07-218).

### sgRNA library design

CRISPRpath sgRNA library was designed to screen cis-regulatory elements for *HPRT1, MSH2, MSH6, MLH1, PMS2* and *PCNA*. ATAC-seq peaks within the region of 1 Mb upstream and 1 Mb downstream of each target gene including TSS and coding regions were selected as targeting regions for the sgRNA library design (**Supplementary Table 1**). We generated a genome-wide sgRNA database containing all the available unique sgRNAs, each followed by a ‘NGG’ PAM sequence. All the designed unique sgRNAs in the target regions were added in the sgRNA library, excluding sgRNAs containing AATAAA, AAAAA, TTTTT or TTTTTT sequences. Unique 20-bp sequences in the target regions that were not followed by the ‘NGG’ or ‘NAG’ PAM sequences were taken as non-targeting control sgRNAs, excluding non-targeting sgRNAs containing TTT, TTNTT, AATAAA, AAAAA, TTTTT or TTTTTT sequences. Then, a guanine nucleotide was added to all the sgRNAs if the sequence did not start with G to increase efficiency of transcription from U6 promoter. Final sgRNA oligos adhered to the following template: 5’-ATATCTTGTGGAAAGGACGAAACACC-[20- or 21-bp sgRNA sequence]-GTTTTAGAGCTAGAAATAGCAAGTTAAAATAAGGC-3’. In total, 35,763 sgRNAs were included in the library (**Supplementary Fig. 1 and Supplementary Table 2**). We retrieved specificity score and off-target site for each sgRNA from GuideScan^18^ (www.guidescan.com) and assigned the specificity score of sgRNAs not existed in the GuideScan database to 0. The high-quality sgRNAs were filtered with specificity score >0.2 and without perfectly matched or 1-2 mismatches off-target sites.

### Oligo synthesis and library cloning

sgRNA library oligos were synthesized by TWIST BIOSCIENCE and amplified with the forward primer 5’-TCGATTTCTTGGCTTTATATATCTTGTGGAAAGGACGAAACAC-3’ and the reverse primer 5’-AACGGACTAGCCTTATTTTAACTTGCTATTTCTAGCTCTAAAAC-3’. We replaced the Cas9 sequence in lentiCRISPR v2 plasmid (Addgene, 52961) with blasticidin S deaminase sequence to construct the lentiCRISPR-v2-Blast-Puro plasmid (Addgene, 167186). The PCR products were purified via gel excision and column purification (Promega, A9282), and then inserted into the BsmBI-digested lentiCRISPR-v2-Blast-Puro vector by Gibson assembly (New England Biolabs, E2621L). The assembled products were transformed into NEB 5-alpha electrocompetent *E. coli* cells (New England Biolabs, C2989K) by electroporation. About 40 million independent bacterial colonies were cultured, and sgRNA library plasmids were extracted with the Qiagen EndoFree Plasmid Mega Kit (Qiagen, 12381). The recovery rate and distribution of the sgRNA library were checked with next generation sequencing (**Supplementary Fig. 2a-d**).

### Lentivirus production and titration

To make the lentiviral library, 5 µg of sgRNA plasmid library was co-transfected with 3 µg of psPAX (Addgene, 12260) and 1 µg of pMD2.G (Addgene, 12259) lentivirus packaging plasmids into 8 million HEK293T cells in a 10-cm dish with PolyJet (SignaGen Laboratories, SL100688). For each individual sgRNA, 3.75 µg of sgRNA plasmid was co-transfected with 2.25 µg of psPAX (Addgene, 12260) and 0.75 µg of pMD2.G (Addgene, 12259) plasmids into 4 million HEK293T cells in a T25 flask with PolyJet (SignaGen Laboratories, SL100688). Media was replaced 12 h after transfection, and harvested every 24 h for a total of three harvests. Harvested media containing the desired virus were filtered through Millex-HV 0.45-µm PVDF filters (Millipore, SLHV033RS) and further concentrated with 100,000 NMWL Amicon Ultra-15 centrifugal filter units (Amicon, UFC910008).

The titer of lentivirus was determined by transducing 500,000 cells with varying amount (0, 0.5, 1.0, 2.0, 4.0 and 8.0 µL) of concentrated virus and polybrene (Millipore, TR-1003-G, 8 µg/mL,). Viral transduction was performed by centrifuging the lentivirus and cell combination at 1000 RCF for 90 min at 37°C. 3 to 4 h later, virus containing media was replaced with fresh media. 24 h after the transduction, transduced cells were dissociated with Accutase, and seeded as duplicates. One replicate was treated with blasticidin (Gibco, A1113903, 4 µg/mL), and the other replicate was not treated with blasticidin. Four days later, the blasticidin resistant cells and control cells were counted to calculate the ratio of infected cells and the viral titer.

### Determining 6TG concentration via killing curve titration

Both CRISPRi and CRISPRn WTC11 iPSCs were used to determine the minimal lethal concentration of 6TG. Cells were seeded in 24 well plates. When the cells reached around 50% confluence (Day 0), they were treated with 6TG concentrations of 0 (control), 20, 40, 60, 80, 100, 120, 140, and 160 ng/mL. Two wells were allocated for each condition. The cells were examined daily and cultured for 7 days. The media was replaced daily with the specified 6TG concentration. After 3 days, wells with 6TG concentration greater than or equal to 100 ng/mL had no surviving cells. On Day 4 of treatment, the wells with 80 ng/mL 6TG treatment had no surviving cells. On the last day of treatment, the wells with 40 and 60 ng/mL treatments had very few surviving cells, while the 20 ng/mL treatment had many surviving cells. Based on these results, we set 80 ng/mL as the minimal lethal concentration for 6TG.

### CRISPRpath screening and sequencing library preparation

CRISPRpath screens were carried out with 72 million doxycycline inducible CRISPRi or CRISPRn iPSCs in biological replicate. The cells for lentiviral transduction were seeded into 6 well plates with 1 million cells per well, and the lentiviral library (MOI = 0.5) was transduced into the iPSCs with 8 µg/mL Polybrene (Millipore, TR-1003-G) and spun at 1000 RCF at 37°C for 90 min. The transduced cells were treated with doxycycline (Sigma, D9891, 2 µM) and blasticidin (Gibco, A1113903, 4 µg/mL) for 4 days. After this doxycycline and blasticidin treatment, 10 million cells were reserved as a control population, and 100 million cells were used for CRISPRpath screen with doxycycline and 6TG (Sigma, A4660) treatment for 7 days. Finally, survival cells were collected from the 6TG treated population.

The genomic DNA was extracted from each sample via cell lysis and digestion (100 mM Tris-HCl pH8.5, 5 mM EDTA, 200 mM NaCl, 0.2% SDS, 100 µg/mL proteinase k), phenol:chloroform (Thermo Scientific, 17908) extraction and isopropanol (Fisher Scientific, BP2618500) precipitation. To amplify the sgRNA sequences from each sample, thirty-two 50 µl PCR reactions were performed using 500 ng genomic DNA for each reaction and NEBNext® High-Fidelity 2X PCR Master Mix (New England Biolabs, M0541S). The purified libraries were sequenced on the NovaSeq 6000 with 150-bp paired-end sequencing. The detail protocol is available on ENCODE portal (https://www.encodeproject.org/documents/2e6451a9-3b98-4d95-922e-a3d8d2100ddf/).

### CRISPRpath data analysis

The sequence files were down-sampled to the same amount of total reads, and then mapped to the sgRNA library with the requirement of exact match of designed sgRNA sequences in the following patten 5’-CCG-[N19 or N20]-GTT-3’. Only the highly specific sgRNAs (specificity score >0.2, without perfectly matched or 1-2 mismatches off-target sites) were used for downstream data analysis. The sgRNA enrichment for each screen was calculated by comparing 6TG treated samples with the associated control samples with edgeR and TMM normalization. We first used edgeR^37^ to calculate *P* value based on negative binomial model for both targeting sgRNAs and non-targeting control sgRNAs. To achieve empirical false discovery rate less than 5%, we then selected a *P* value cutoff corresponding to the 5% percentile of *P* values from non-targeting control sgRNAs. Finally, we defined enriched sgRNAs with *P* value less than the selected *P* value cutoff, and fold change >2. The ATAC-seq peaks were identified as functional enhancers for the six MMR genes by having at least 3 significant enriched sgRNAs. Analysis scripts are available at https://github.com/MichaelMW/crispy.

### Analysis of genomic feature and chromatin signature of identified enhancers

Genomic distances between enhancer and TSS pairs were calculated based on the distance from the center of enhancers to the transcription start sites of the target genes. The number of interval genes is the number of all the RefSeq annotated genes between each enhancer and paired target gene. The signal of chromatin signatures, including ATAC-seq, H3K27ac, H3K4me3, CTCF binding and RNA-seq, were calculated by deeptools (v3.4.3)^38^. The enhancer-like promoters are the enhancers overlap with the region 500 bp upstream and downstream of a RefSeq annotated TSS.

### Validation of identified enhancers using CRISPRi

We cloned lentiCRISPR-v2-HygR-EGFP (Addgene, 167188) and lentiCRISPR-v2-HygR-mCherry (Addgene, 167189) vectors by replacing the Cas9 and puromycin N-acetyltransferase sequences in lentiCRISPR v2 plasmid (Addgene, 52961) with hygromycin B phosphotransferase and EGFP or mCherry sequences. To validate the identified enhancers, individual sgRNAs targeting identified enhancers were cloned into the lentiCRISPR-v2-HygR-GFP or lentiCRISPR-v2-HygR-mCherry vector. The doxycycline inducible CRISPRi WTC11 iPSCs were infected with the lentivirus expressing sgRNAs for three replicates per sgRNA. The sgRNA infected cells were grown with hygromycin (Gibco, 10687010, 150 µg/mL) and doxycycline (Sigma, D9891, 2 µM) containing media. Seven days later, the cells were collected and total RNA was extracted from the cells using the Qiagen RNeasy® Plus Kit (Qiagen, 74134). One µg of RNA was then used to synthesize cDNA using the Bio-RAD iScript cDNA Synthesis Kit (Bio-RAD, 1708840). qPCR reactions for targeted genes were performed with the Luminaris HiGreen qPCR Master Mix (Thermo Scientific, K0993) on the Roche LightCycler 96 System. The qPCR primers are listed in **Supplementary Table 4** and the sgRNA sequences are listed in **Supplementary Table 6**. For each tested element in **Fig. 3a-c** and **Supplementary Fig. 6b**, we performed CRISPRi experiments with two independent sgRNAs and used the results from the sgRNA with stronger transcriptional repression in **Fig. 4b**.

### shRNA mediated RNA interference

shRNAs were designed by using DSIR tool (http://biodev.extra.cea.fr/DSIR/DSIR.html) targeting *SOCS5, FOXN2* and *TMEM230*. The sequences of shRNAs are listed in **Supplementary Table 5**. The shRNAs were cloned into lentiCRISPR-v2-HygR-mCherry vector under the control of human U6 promoter and packaged into lentivirus for cell transduction. The WTC11 iPSCs transduced with shRNA lentivirus were treated with hygromycin (Gibco, 10687010, 150 µg/mL) for 7 days and then collected for RNA extraction and qPCR.

### ATAC-seq

ATAC-seq was carried out using the Nextera DNA Library Prep Kit (Illumina, FC-121-1030) as previously described^39^. The detailed protocol is available on the ENCODE portal (https://www.encodeproject.org/documents/0317894c-5a42-4f03-b865-c2a2d08708ef/). Briefly, each library started with 100,000 fresh iPSCs, and the cells were incubated with ice cold nuclei extraction buffer (10 mM TrisHCl pH 7.5, 10 mM NaCl, 3 mM MgCl2, 0.1% Igepal CA630, and 1x protease inhibitor) for 5 min on ice, then centrifuged at 500 RCF for 5 min. 50,000 resulting nuclei were treated with tagmentation buffer (25 µL Buffer TD with 50,000 nuclei, 22.5 µL water, 2.5 µL TDE1) for 30 min at 37°C. The transposed DNA was purified using Qiagen MinElute PCR purification kit (Qiagen, 28006), amplified using Nextera primers, then size-selected for fragments between 150 and 1000 bp using SPRISelect beads (Beckman Coulter, B23319). Libraries were sent for single-end sequencing on the HiSeq 4000 (50 bp single-end reads). Reads were mapped to hg38/GRCh38 and processed using the ENCODE pipeline (https://github.com/kundajelab/atac_dnase_pipelines, V1.8.0), which ran on the default settings. The ATAC-seq peaks were filtered with FDR cutoff of 0.1%, and adjacent peaks were merged if they are less than 1 kb apart.

### RNA-seq

RNA was extracted from fresh cells using the RNeasy Plus Mini Kit (Qiagen, 74134). Approximately 1000 ng of extracted RNA was used to prepare libraries for sequencing using the TruSeq Stranded mRNA Library Prep Kit (Illumina, 20020594). Libraries were sent for paired-end sequencing on the NovaSeq 6000 (100 bp paired-end reads). Reads were aligned to hg38/GRCh38 using STAR 2.7.0f^40^ with the standard ENCODE settings, and transcript quantification was performed in a strand-specific manner using RSEM 1.3.1^41^ with the annotation from GENCODE v32. Only the first read was used, and all reads were trimmed to 51bp using TrimGalore 0.4.5 running the following options: -q 20 --length 20 -- stringency 3 -- trim-n. The edgeR package in R (3.20.9)^37^ was used to calculate TMM-normalized FPKM values for each gene based on the expected counts and gene lengths for each library. The mean gene expression across all replicates was used for analysis.

### ChIP-seq

ChIP-seq libraries were constructed from 2 million WTC11 iPSCs. Cells were crosslinked in 1% formaldehyde at room temperature for 20 min and then quenched with 2.5 M glycine at room temperature for 5 min. Fixed cells were lysed and chromatin was sonicated by Covaris with the following parameters: Duty Factor: 2%, Peak Incident Power: 105W, Cycles per Burst: 200, for 30 min. Input chromatin was removed and stored at -20°C for later processing. Magnetic beads (Invitrogen, Dynabeads Protein A, 10001D) were preincubated with H3K27ac antibody (Active Motif, 39133, Lot 22618011) for 2 hours at 4°C before being added to sheared chromatin. Samples were incubated overnight at 4°C. Beads were washed 3 times and chromatin was then eluted. Samples were incubated at 65°C overnight to reverse the crosslinking. DNA was treated with RNase A for 1 hr at 37°C and Proteinase K (New England Biolabs, 8107) for 1 hr at 55°C. DNA was purified by phenol-chloroform extraction and ethanol precipitation. Libraries were prepared using Tru-seq adapters and size-selected using SPRIselect beads prior to amplification and paired-end sequencing. Libraries were sent for paired-end sequencing on the NovaSeq 6000 (150 bp paired-end reads). Sequencing reads were trimmed to 50 bp and mapped to hg38 using bowtie2 with the following options: --local -- very-sensitive-local --no-unal --no-mixed --no-discordant --phred33 -I 10 -X 700. Picard Tools was used to remove blacklisted regions and duplicate reads and MACS2 was used to call peaks on merged replicates at an FDR cutoff of 1%.

### CUT&Tag

CUT&Tag libraries were constructed from 150,000 WTC11 iPSCs according to previously described methods^42^. Cells were lysed in nuclei extraction buffer (20 mM HEPES-KOH pH 7.9, 10 mM MgCl2, 0.1% Triton X-100, 20% glycerol and 1x protease inhibitor) on ice for 10 min. The samples were spun and resuspended in 100 µl nuclei extraction buffer. Meanwhile, 10 µl of BioMag Plus Concanavalin A (Bangs Laboratories, BP531) were equilibrated in binding buffer (1x PBS, 1 mM CaCl_2_, 1 mM MgCl_2_ and 1 mM MnCl_2_). The equilibrated beads were added to the samples and incubated with rotation for 15 min at 4°C. Nuclei-bound beads were washed with Buffer 1 (20 mM HEPES-KOH pH 7.9, 150 mM NaCl, 2 mM EDTA, 0.5 mM spermidine, 0.1% BSA and 1x protease inhibitor) and Buffer 2 (20 mM HEPES-KOH pH 7.9, 150 mM NaCl, 0.5 mM spermidine, 0.1% BSA and 1x protease inhibitor). After washing nuclei-bound beads were resuspended in 50 µl Buffer 2 with 0.5 µl antibody (H3K4me3 from Millipore, 04-745, Lot 3543820 and CTCF from Millipore, 07-729, Lot 3059608) and incubated with rotation overnight at 4°C. Samples were washed twice with Buffer 2 and resuspended in 50 µl Buffer 2 with antibody (antibodies-online Inc., Guinea Pig anti-Rabbit IgG, ABIN101961, Lot 42323) and incubated for 1 hr at room temperature with rotation. Samples were washed again with Buffer 2 and resuspended in 100 µl Buffer 3 (20 mM HEPES-KOH pH 7.9, 300 mM NaCl, 0.5 mM Spermidine, 0.1% BSA and 1x proteinase inhibitor) containing 0.04 µM pA-Tn5. Samples were incubated for 1 hr at room temperature, washed three times with Buffer 3 and resuspended in tagmentation buffer (20 mM HEPES-KOH pH 7.9, 300 mM NaCl, 0.5 mM Spermidine, 10 mM MgCl_2_, 0.1% BSA and 1x proteinase inhibitor). Samples were incubated for 1 hr at 37°C. Samples were treated with Proteinase K (New England Biolabs, 8107) for 1 hr at 50°C. DNA was purified by phenol-chloroform extraction and ethanol precipitation. Libraries were prepared using Tru-seq adapters and size-selected using SPRIselect beads prior to amplification and paired-end sequencing. Libraries were sent for paired-end sequencing on the Mini-seq (37 bp paired-end reads, H3K4me3 libraries) or NovaSeq 6000 (150 bp paired-end reads, CTCF libraries). Sequencing reads (CTCF libraries were trimmed to 50 bp) were mapped to hg38 using bowtie2 with the following options: --local --very-sensitive-local --no-unal --no-mixed --no-discordant --phred33 -I 10 -X 700. Picard Tools was used to remove blacklisted regions and duplicate reads and SEACR^43^ was used to call peaks on merged replicates.

### H3K4me3 PLAC-seq

H3K4me3 PLAC-seq data in WTC11 cells were generated as previously described^44^ in biological replicates (clone 6 and clone 28) (https://data.4dnucleome.org/experiment-set-replicates/4DNESDRL4ZKM/ and https://data.4dnucleome.org/experiment-set-replicates/4DNESIZ5TTHO/). We combined the two biological replicates, and applied the MAPS pipeline^45^ to identify significant long-range chromatin interactions at 5 kb bin resolution for the genomic distance 10 kb ∼ 1 Mb. The reference genome is GRCh38. In addition, for each 5 kb bin pair anchored at H3K4me3 peaks, the MAPS pipeline outputs the normalized contact frequency, which adjusts for the biases from effective fragment length, GC content, sequence mappability, H3K4me3 enrichment level and 1D genomic distance effect.

### Comparison between strong enhancers and weak enhancers using H3K4me3 PLAC-seq data

For 4 genes *HPRT1, MLH1, PMS2* and *PCNA*, there are 23 enhancer-promoter pairs between strong enhancers and their target genes, and 21 enhancer-promoter pairs between weak enhancers and their target genes. We mapped each enhancer and promoter of target gene into 5 kb bins, and obtained the distance normalized H3K4me3 PLAC-seq contact frequency for 5 kb bin pairs containing the enhancer-promoter pairs. Since *MSH2* and *MSH6* are located within 407 kb linear genomic distance with each other and we can’t assign enhancers to either gene reliably, enhancers identified near *MSH2* and *MSH6* were excluded from this analysis.

### Comparison between enhancer-like promoters and control promoters using H3K4me3 PLAC-seq data

For this analysis, control promoters are active promoter regions with annotated ATAC-seq peaks and tested negative as enhancers for the for MMR genes. We mapped each promoter into a 5 kb bin that was used in the PLAC-seq analysis. We only choose the bins with one annotated active promoter, which gave us 31 enhancer-like promoters and 43 control promoters in this analysis. We counted the number of significant H3K4me3 PLAC-seq interactions anchored at the 5 kb bins with these promoter sequences. In addition, as described in our previous study^23^, for promoters with at least one significant interaction, we calculated the summation of –log10 FDR of significant interactions, which is a measure of the overall interaction strength.

### Chromatin contact frequency comparison between strong enhancers and weak enhancers in K562 cells and mESCs

For the chromatin contact frequency comparison of enhancers in K562 cells and mouse embryonic stem cells (mESCs), we downloaded the identified enhancers from each publication^6, 8^, and defined strong enhancer with cutoff of 50% ≤ transcriptional contribution ≤ 100%, weak enhancer with cutoff of 0% < transcriptional contribution ≤ 20%. H3K27ac HiChIP data in K562 cells^46^ and H3K4me3 PLAC-seq data in mESCs^45^ were used for comparison. The comparisons were performed in 10 kb resolution.

### Motif scan and Transcription factor identification

The fasta files were first generated in the hg38 genome for the identified strong enhancers and weak enhancers separately. For each strong enhancer and weak enhancer, the FIMO software (version 5.1.0)^47^ with human motif database HOCOMOCO (v11 FULL)^48^ was used to scan the motifs. All the FIMO motif scans were in default settings. We then filtered the transcription factors (TFs) in each strong and weak enhancer loci by FDR cutoff of 0.05 and p-value cutoff of 0.0001 and gene expressions cutoff of FPKM > 1. By taking the TFs with TF motif appeared in more than 80% enhancers, 47 TFs were considered as commonly appearing in the strong enhancers, and 35 TFs were in the weak enhancers.

## Acknowledgements

This work was supported by the National Institutes of Health grants UM1HG009402 (to Y.S., B.R., and W.W.) and U54DK107977 (to B.R. and M.H.).

## Data availability

The CRISPRpath screen datasets used in this study are available at the ENCODE portal (www.encodeproject.org, accession number: ENCSR617AZY (sgRNA plasmid library), ENCSR427OPP (CRISPRi control), ENCSR900AXT (CRISPRi 1X), ENCSR254SJU (CRISPRi 2X), ENCSR793DSE (CRISPRi 3X), ENCSR250ZWC (CRISPRn control), ENCSR117YGQ (CRISPRn 1X), ENCSR071ZGB (CRISPRn 2X), ENCSR482PHH (CRISPRn 3X)). WTC11 iPSCs H3K4me3 PLAC-seq datasets are available at the 4DN data portal (data.4dnucleome.org, accession number: 4DNESIZ5TTHO and 4DNESDRL4ZKM). ATAC-seq, ChIP-seq, CUT&Tag, and RNA-seq datasets in WTC11 iPSCs are available at the Gene Expression Omnibus under the accession number GSE166839. Data can be visualized on the WashU Epigenome Browser using the session at the following link (https://epigenomegateway.wustl.edu/browser/?genome=hg38&sessionFile=https://shen-xren.s3-us-west-1.amazonaws.com/CRISPRpath/eg-session-QRXJ0218-4d710b60-6ea7-11eb-8d8d-03c7189570c0.json). Tracks include ATAC-seq, H3K27ac, H3K4me3, and CTCF signals, and the identified enhancers from CRISPRi 2X and 3X screens. RefGene 38 genes are also displayed. The plasmids generated in this study are available from Addgene (#167186, #167188, #167189).

## Code availability

The computer code used for analyzing CRISPRpath datasets is available at https://github.com/MichaelMW/crispy.

## Author contributions

X.R. and Y.S. designed the study. X.R. and B.L. designed the sgRNA library under the supervision of Y.S. and B.R. X.R., K.J., I.R.J., M.A.T., J.L., L.M., and T.W.T. performed the experiments. M.W. designed the CRISPY under the supervision of W.W. X.R., B.L., L.Z., G.L., and Y.L. performed data analysis. M.Y. and R.H. constructed the H3K4me3 PLAC-seq libraries. L.L., A.A. and M.H. analyzed PLAC-seq and HiChIP data. X.R. and Y.S. prepared the manuscript with input from all other authors.

---

**Supplementary Table 1. ATAC-seq peak regions used to design the sgRNA library**.

**Supplementary Table 2. List of sgRNA sequences used for CRISPRpath screen**.

**Supplementary Table 3. Enhancers identified from CRISPRn 2X, CRISPRi 2X and CRISPRi 3X screens**.

**Supplementary Table 4. List of primers used for RT-qPCR**.

**Supplementary Table 5. List of shRNA sequences used for RNA interference experiments**.

**Supplementary Table 6. List of sgRNA sequences used for CRISPRi-mediated enhancer validation experiments**.

**Supplementary Table 7. Pairwise comparisons of data in Figure 4c**.

**Supplementary Figure 1.**
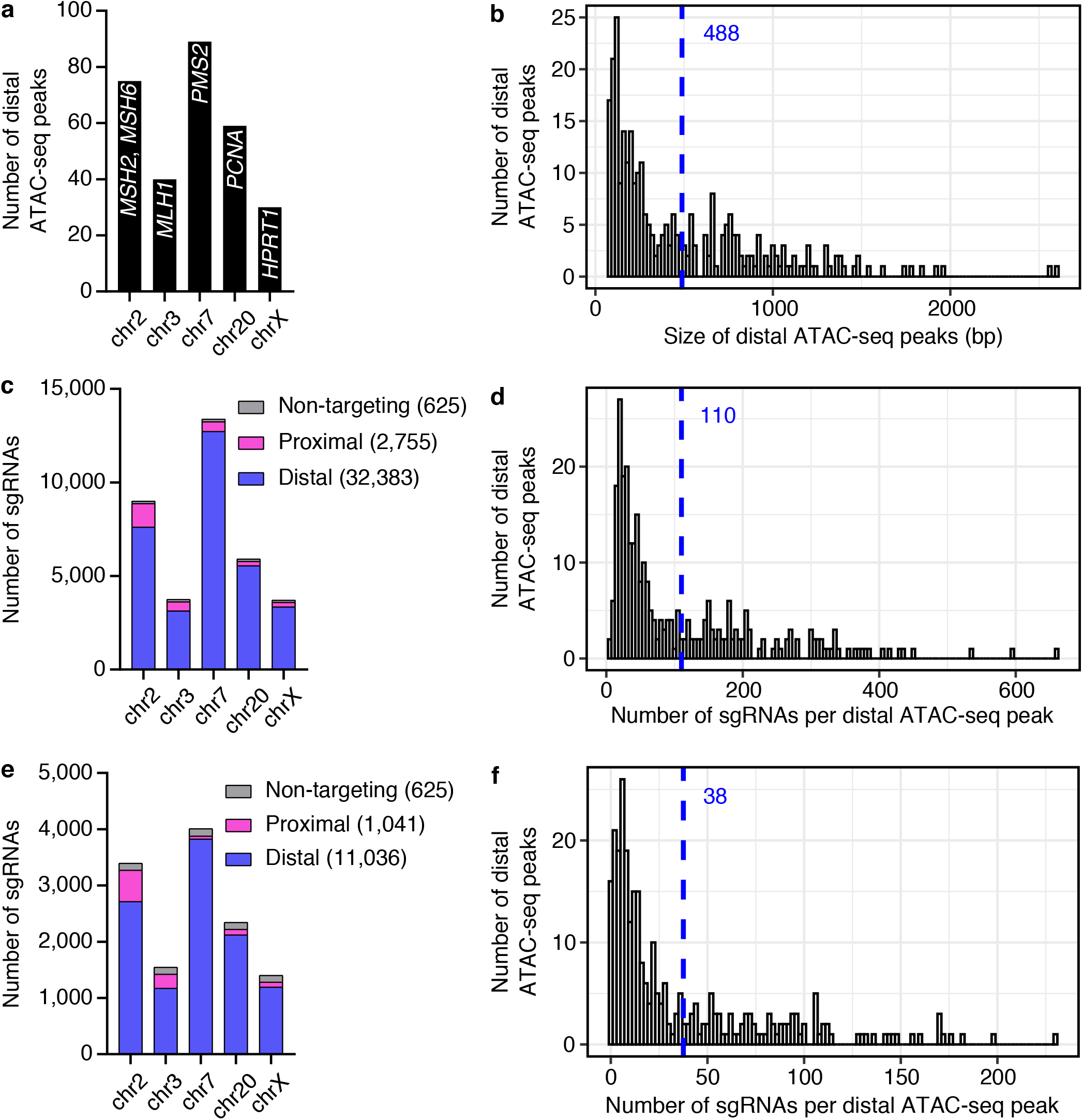
Features of sgRNA library for CRISPRpath screen. (**a**) Bar graph shows the number of distal ATAC-seq peaks used as candidate CREs for six target genes. (**b**) Histogram shows size distribution of distal ATAC-seq peaks. The average size is 488 bp (blue dash line). (**c, e**) The composition of the sgRNA library. In total, 35,763 sgRNAs were included in the library (**c**), and 12,702 sgRNAs are high quality sgRNAs (**e**). (**d, f**) Distribution of the number of sgRNAs per distal ATAC-seq peak. Average numbers of sgRNA per ATAC-seq peak are indicated with blue dash lines.

**Supplementary Figure 2.**
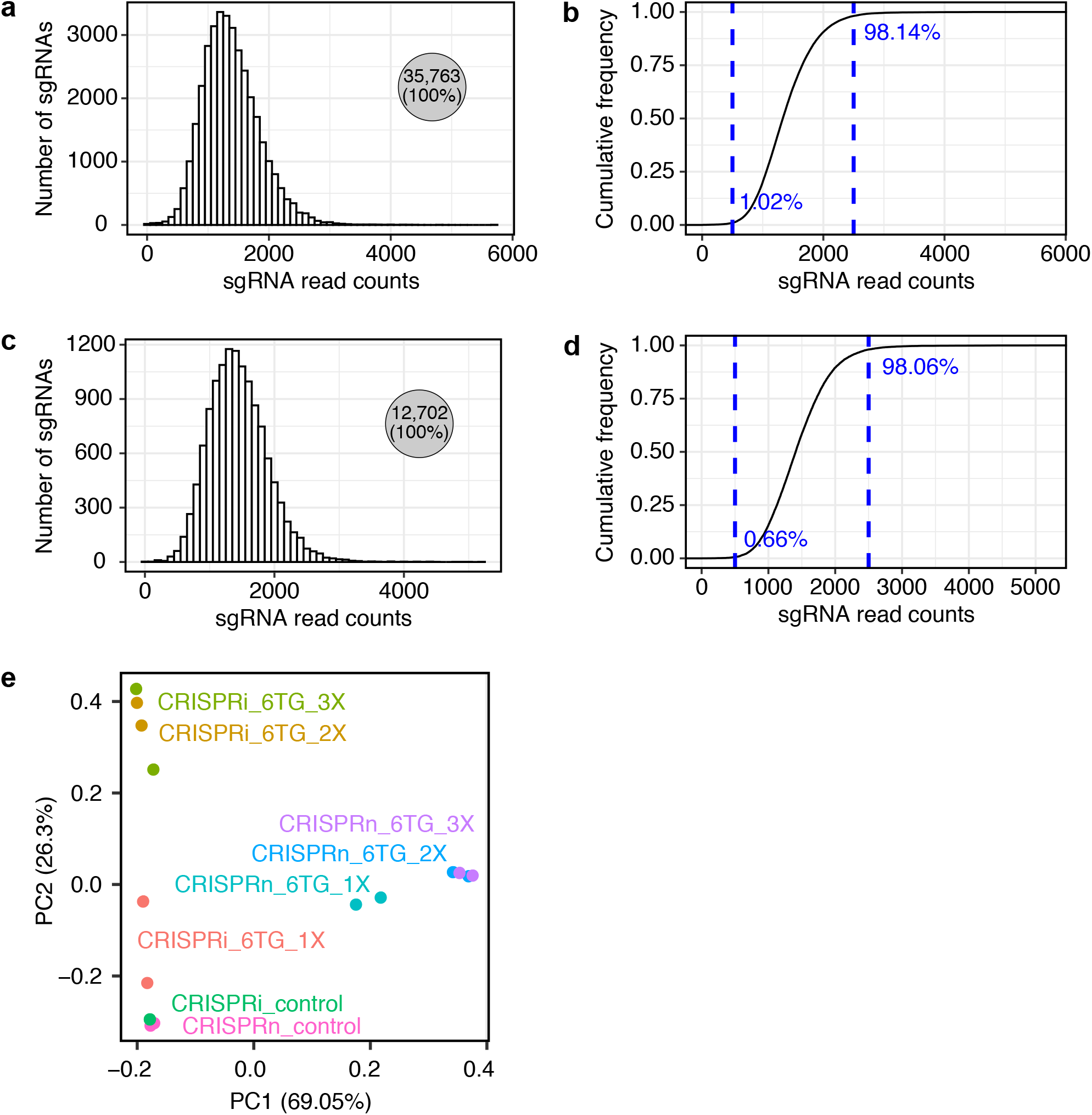
Quality of the sgRNA library and CRISPRpath screen libraries. (**a**) Distribution of sgRNA oligo read counts in the sgRNA library. (**b**) Cumulative frequency of sgRNAs in the sgRNA library. (**c**) Distribution of high quality sgRNAs read counts in the sgRNA library. (**d**) Cumulative frequency of high quality sgRNAs in the sgRNA library. The constructed sgRNA plasmid library recorvered all the designed sgRNAs with the copy number difference less than five fold for at least 97% designed sgRNAs. (**e**) PCA analysis shows the high reproducibility of the CRISPRpath screen libraries between biological replicates.

**Supplementary Figure 3.**
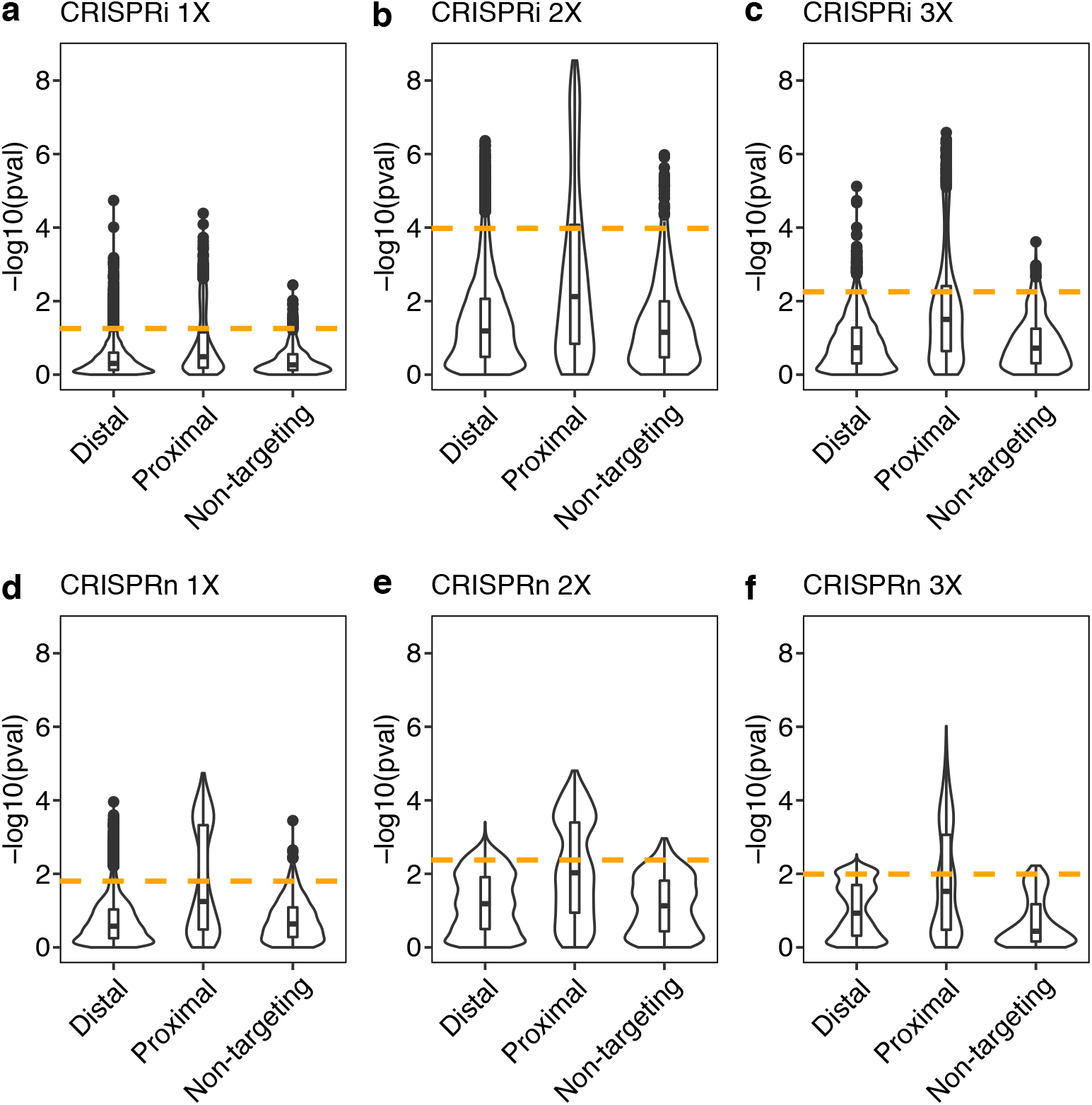
P value cutoff used for identifying enriched sgRNAs from each screens. (**a-f**) Distribution of *P* value for tested distal, proximal and non-targeting control sgRNA groups. Orange dash lines indicate 5% percentile of the *P* values from non-targeting control sgRNAs to achieve a false discovery rate of 5%.

**Supplementary Figure 4.**
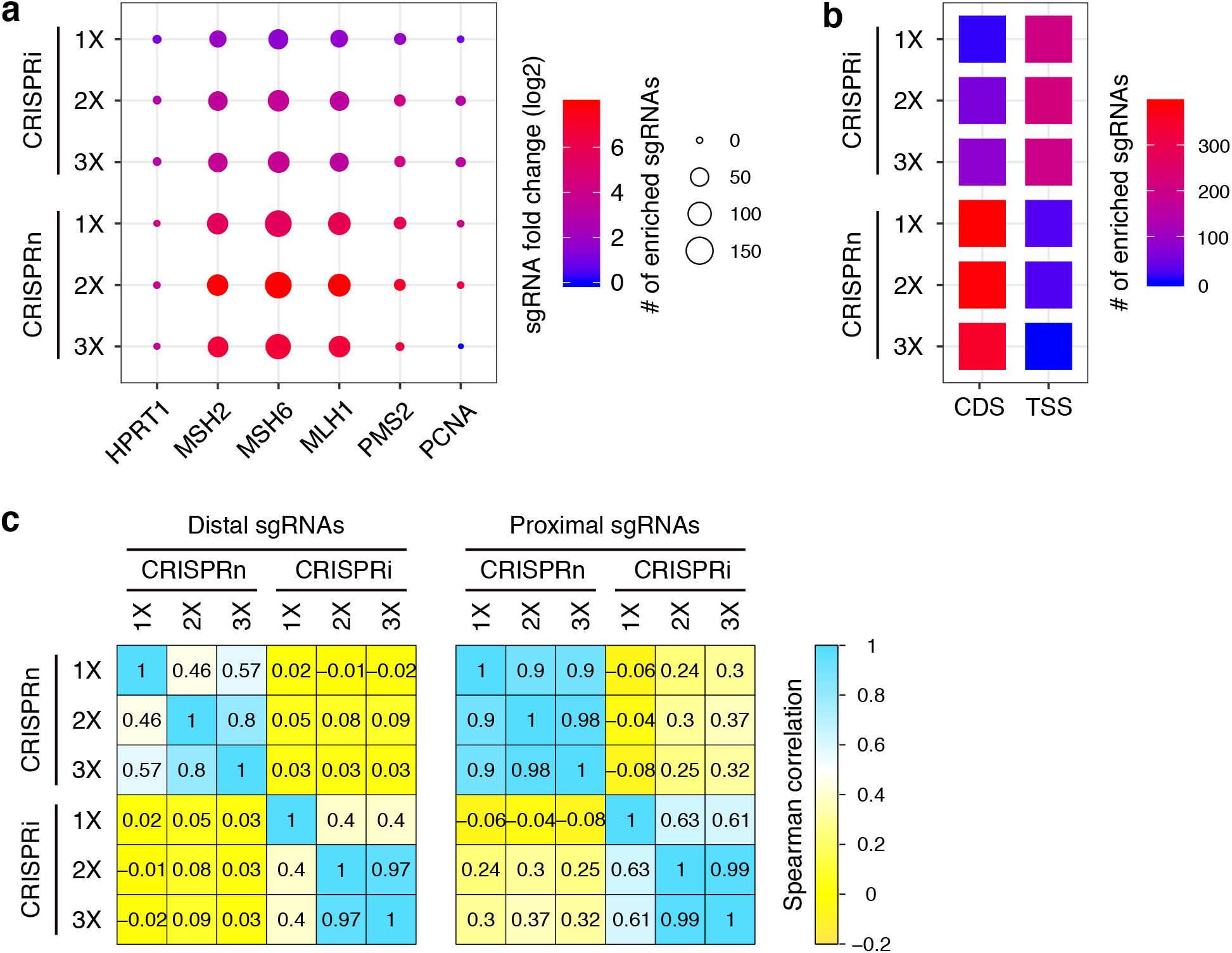
Enriched proximal sgRNAs and sgRNA ranking analysis. (**a**) Number and fold change of the enriched proximal sgRNAs for the six target genes from CRISPRi and CRISPRn screens. The color indicates fold changes, and the size of circle indicates the number of enriched sgRNAs. (**b**) Enrichment analysis shows the enriched proximal sgRNAs bias towards to the TSS region for CRISPRi screens and the protein coding region (CDS) for CRISPRn screens. Color represents the number of enriched sgRNAs. (**c**) Spearman correlation analysis of the distal and proximal sgRNAs ranking shows proximal sgRNAs exhibiting higher correlation between each screen compared to distal sgRNAs.

**Supplementary Figure 5.**
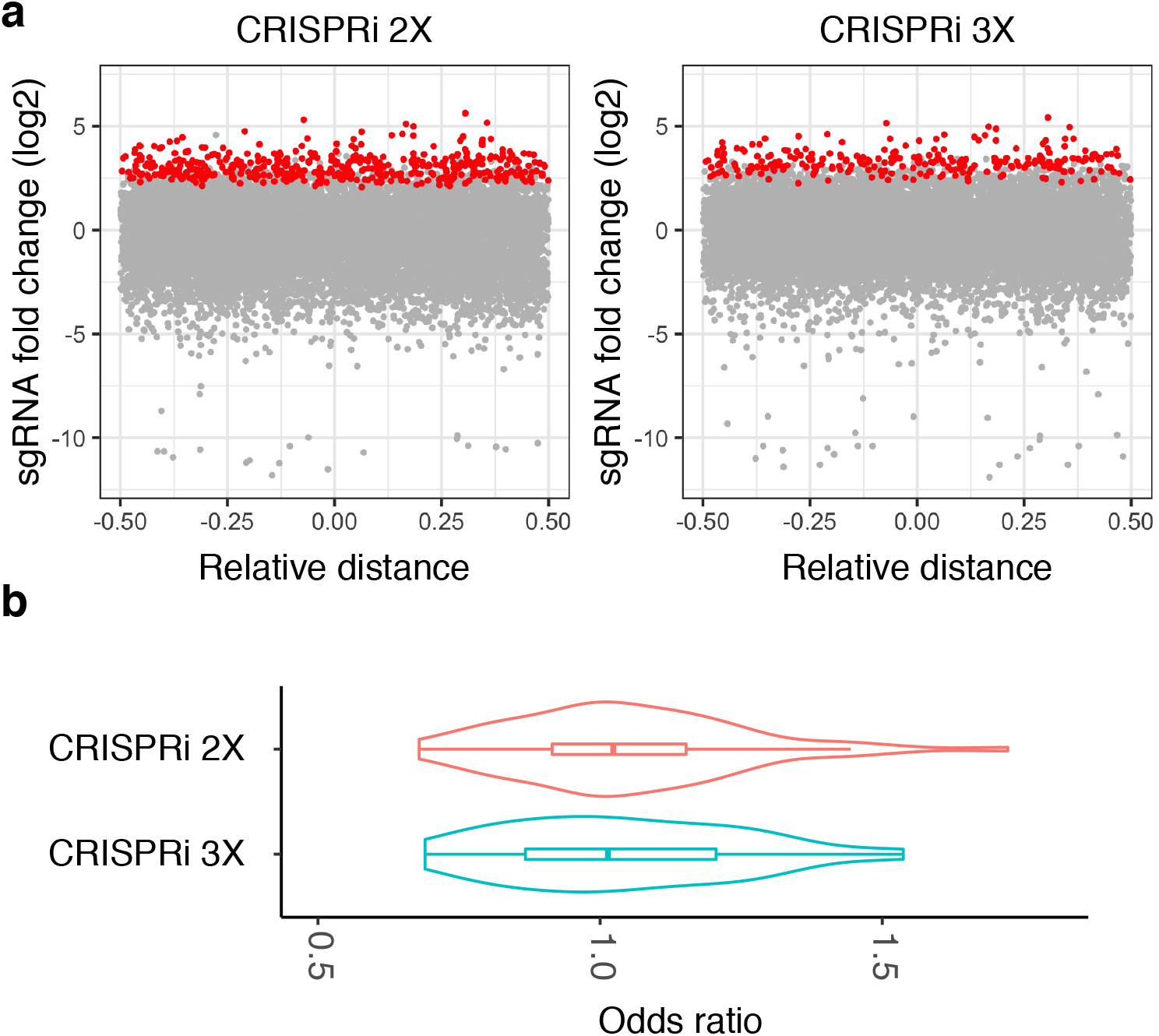
Enriched sgRNAs identified from CRISPRi screens exhibit no position and strand preference. (**a**) Enriched sgRNAs from CRISPRi 2X (red dots, n=448) and CRISPRi 3X (red dots, n=260) screens showed similar distributions across candidate CREs. (**b**) Odds ratio analysis of the fold change of enriched sgRNAs shows enriched sgRNAs have no strand preference. Enhancers with enriched sgRNAs only targeting one strand were exclude for the analysis. Odds ratio was calculated for each element with the equation of ave(log_2_(fold change of sgRNA targeting plus strand)) / ave(log_2_(fold change of sgRNA targeting minus strand)). Violin plots show the distributions of odds ratio values within each screen, and boxplots indicate the median, IQR, Q1 − 1.5 × IQR and Q3 + 1.5 × IQR.

**Supplementary Figure 6.**
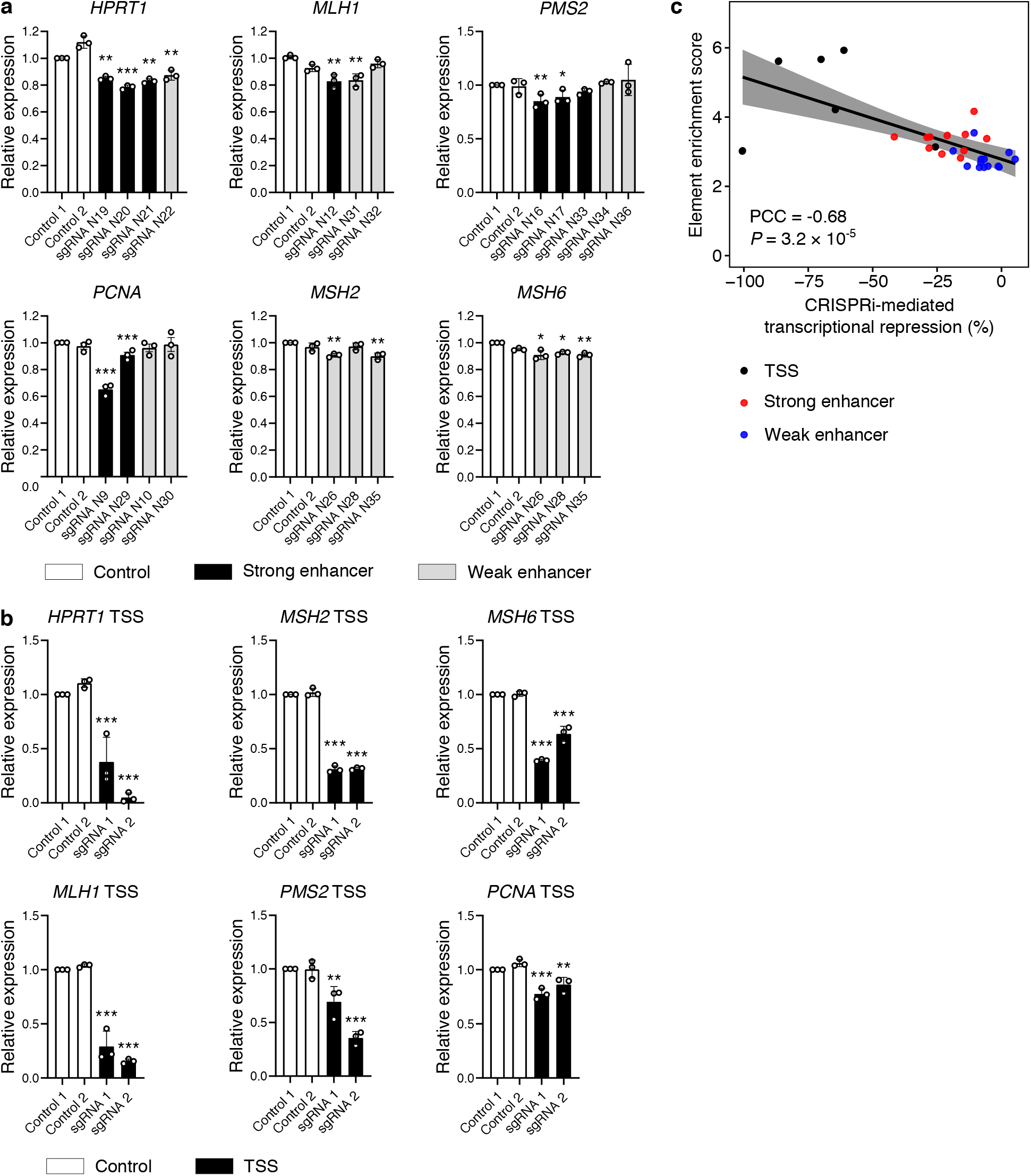
Validation of CIRSPRpath identified enhancers. (**a**) Validation of the strong (black) and weak (grey) enhancers with CRISPRi followed by RT-qPCR. Three independent replicates per condition. The significance was calculated with two-tailed two-sample *t*-test. Data are mean and s.d. * *P* < 0.05, ** *P* < 0.01, *** *P* < 0.001. (**b**) CRISPRi-mediated transcriptional repression of six target genes by targeting TSS of each gene. Three independent replicates per condition. The significance was calculated with two-tailed two-sample *t*-test. Data are mean and s.d. * *P* < 0.05, ** *P* < 0.01, *** *P* < 0.001. (**c**) Pearson correlation analysis reveals element enrichment score from CRISPRpath screens correlates with element effect size on transcription from CRISPRi (Pearson correlation, PCC = -0.68, *P* = 3.2 × 10^−5^).

**Supplementary Figure 7.**
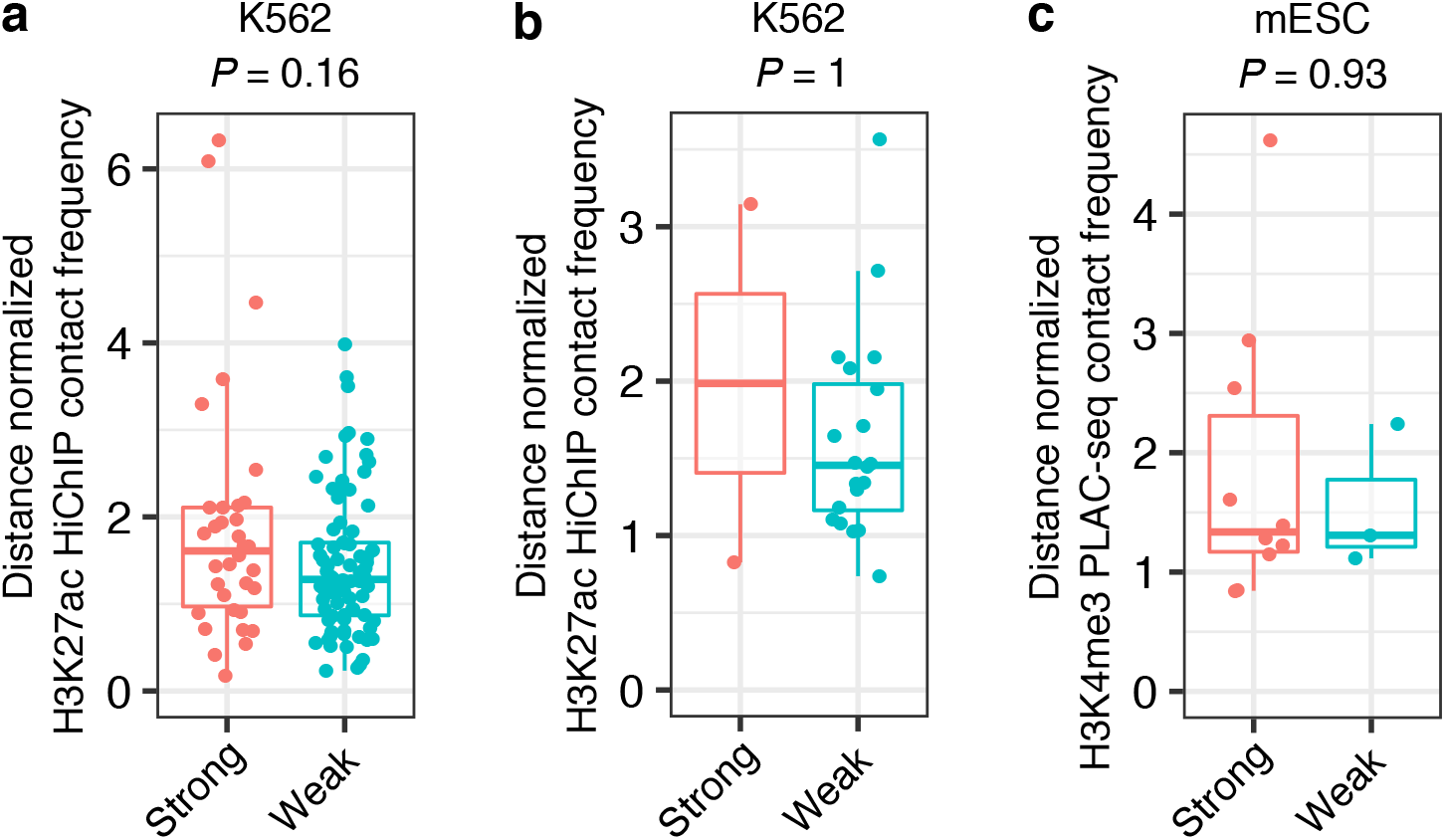
Chromatin contract frequency analysis for the enhancers in K562 cells and mESCs. (**a**) Distance normalized H3K27ac HiChIP contact frequency for strong (n = 34) and weak (n = 82) enhancers identified with crisprQTL mapping in K562 cells. (**b**) Distance normalized H3K27ac HiChIP contact frequency for strong (n = 2) and weak (n = 20) enhancers identified with CRISPRi-FlowFISH screen in K562 cells. (**c**) Distance normalized H3K4me3 PLAC-seq contact frequency for strong (n = 10) and weak (n = 3) enhancers identified in mouse embryonic stem cells. Boxplots indicate the median, IQR, Q1 − 1.5 × IQR and Q3 + 1.5 × IQR. *P* values are from Wilcoxon test.

